# Sensitizing the Cell Killing Effects of Doxorubicin as a Possible Therapeutic Strategy Against Glioblastoma Multiforme

**DOI:** 10.64898/2025.12.23.696220

**Authors:** Anthony Berdis

## Abstract

Glioblastoma multiforme (GBM), the most common type of brain cancer, is also the deadliest of all cancers, having 5-year survival rates of less than 10%. Standard therapy for GBM includes surgery followed by ionizing radiation and treatment with temozolomide, a DNA alkylating agent. Unfortunately, even with aggressive treatments with these modalities, the median survival time for most GBM patients is less than 16 months. This study examines a new treatment strategy by combining doxorubicin, a soluble DNA-damaging agent that produces double-stranded DNA like ionizing radiation, with an artificial nucleoside designated 5-nitroindolyl-2’deoxyriboside (5-NIdR) that inhibits the misreplication of DNA lesions produced by the drug. Cell-based studies with grade 3 and grade 4 astrocytomas demonstrate that combining 5-NIdR with doxorubicin improves the cell-killing effects of the drug by increasing apoptosis. Flow cytometry experiments demonstrate that 5-NIdR does not increase the level of reactive oxygen species produced by doxorubicin. Instead, the increase in cell death produced by combining the two agents likely reflects inhibiting the replication of damaged DNA. Consistent with this mechanism, cell-cycle experiments demonstrate that this combination blocks cells at G_o_/G_1_, suggesting that 5-NIdR interferes with DNA polymerase activity during the repair of double-strand DNA breaks. The ability of 5-NIdR to inhibit double-strand DNA break repair could be extended to other modalities used to treat brain cancer such as ionizing radiation. Furthermore, the chemosensitizing effect of 5-NIdR could improve the overall efficacy of doxorubicin by lowering the risk of side effects caused by the DNA-damaging agent.

## Introduction

Glioblastoma multiforme (GBM) is the most common type of brain cancer, afflicting over 12,000 children and adults each year in the United States [1]. GBM is also the deadliest of all cancers, having 5-year survival rates of less than 10% [2]. Standard therapy for GBM includes surgery, ionizing radiation (IR) and treatment with the DNA-alkylating agent, temozolomide (TMZ). Unfortunately, even with aggressive treatments with IR and TMZ, the median survival time for most GBM patients is less than 16 months [3].

At the molecular level, TMZ and IR produce DNA lesions that can be misreplicated to promote the generation of genetic mutations that can drive drug resistance and produce more aggressive malignancies. TMZ generates several DNA lesions including N^3^-methyladenine, O^6^-methylguanine, and N^7^-methylguanine. Each structurally distinct form of DNA damage can produce cytostatic and cytotoxic effects [4]. However, N^7^-methylguanine is perhaps the most important since spontaneous depurination of the methylated base creates a highly toxic DNA lesion known as an abasic site. [5]. Exposure to IR also produces a wide range of DNA lesions. However, the most cytotoxic is a double strand DNA break (DSB) which is a non-templating DNA lesion similar to an abasic site. In both cases, the presence of non-templating DNA lesions can pose unique challenges to a cancer cell – either stop synthesizing DNA and die or replicate the DNA lesion to potentially introduce genomic errors. Unfortunately, most cancers chose the latter.

The replication of damaged DNA is a process termed translesion DNA synthesis (TLS) [6–8]. This process allows damaged DNA to be effectively by-passed, providing a simple pathway allowing cancer cells to survive the cell-killing effects of certain chemotherapeutic agents (Figure 1A). As such, TLS activity can partially be responsible for the onset of resistance to many DNA-damaging agents [9,10]. In addition, TLS activity is often pro-mutagenic and can generate more mutations in treated cancer cells. Both features produce negative consequences for patients as increased drug resistance and induced mutagenesis can work in tandem to generate more aggressive malignancies.

**Figure 1.**
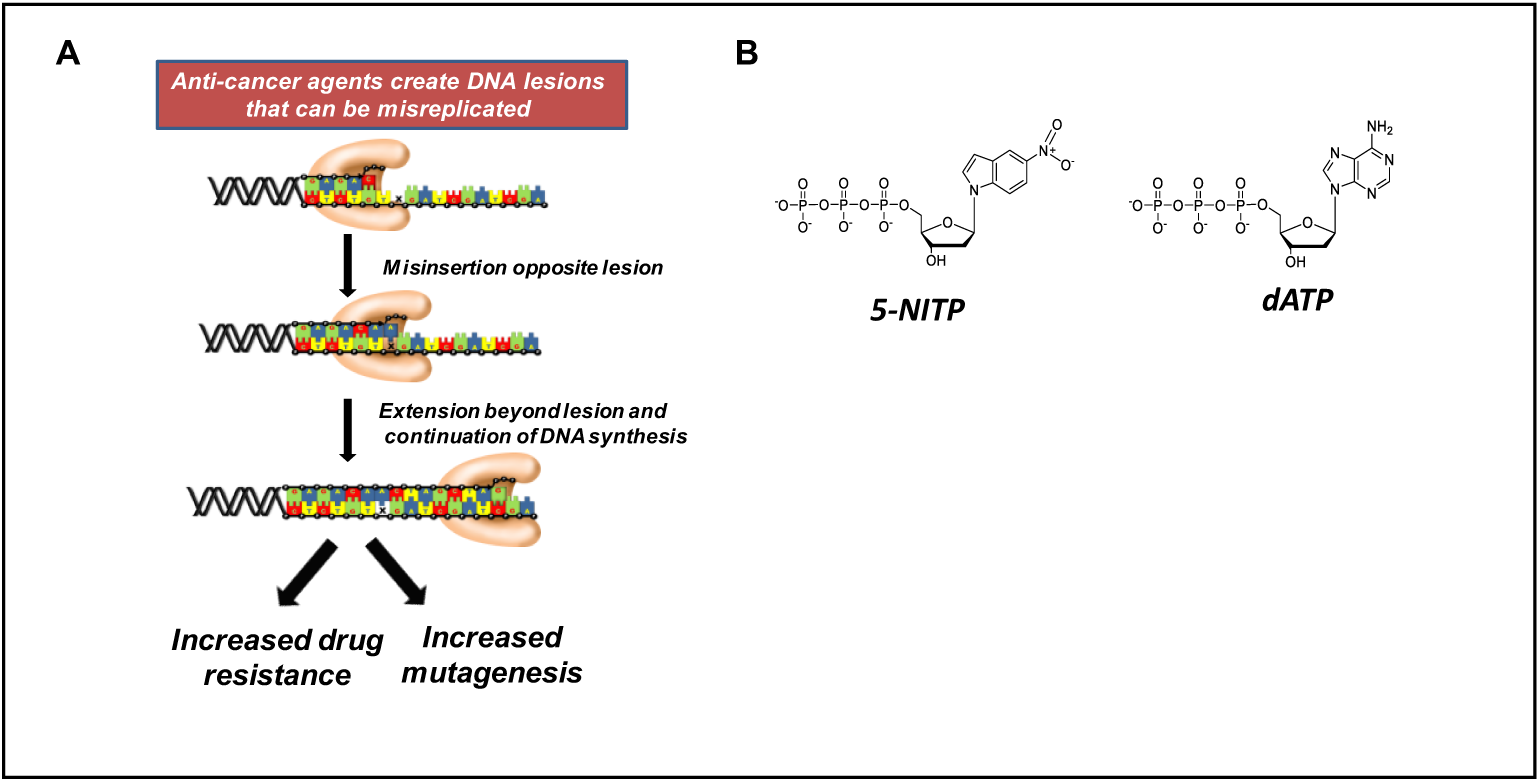
(A) Model for drug resistance caused by DNA polymerases misreplicating DNA lesions generated by anti-cancer agents that damage DNA. In addition to driving drug resistance, the misreplication of damaged DNA can increase mutagenesis and induce genetic drift within a population of tumor cells. (B) Structural comparison between the artificial deoxynucleotide, 5-NITP, and the natural purine nucleoside triphosphate, dATP.

To combat these problems, we developed an artificial nucleotide analog designated **<u>5</u>**-***<u>N</u>***itroIndolyl ***<u>T</u>***ri***<u>P</u>***hosphate (5-NITP) (Figure 1B) that inhibits TLS activity *in vitro* and *ex vivo* models of cancer [11–15]. At the molecular level, 5-NITP mimics the shape and size of dATP but lacks conventional hydrogen bonding groups necessary for normal DNA replication. Thus, the artificial nucleotide is poorly incorporated opposite undamaged DNA and cannot efficiently inhibit normal DNA synthesis [11]. Instead, the lack of hydrogen bonding groups allows the analog to be efficiently and selectively incorporated opposite non-templating DNA lesions such as those produced by TMZ [12, 13]. Once inserted opposite a DNA lesion, the analog is refractory to elongation and terminates pro-mutagenic DNA synthesis [14]. The inhibitory effect on TLS ultimately disrupts the continuity of chromosomal to induce cell death [11–15].

This report assesses the ability of 5-NIdR to potentiate the cell-killing effects of another DNA damaging, doxorubicin (DOX), against different GBM cell lines. DOX was chosen since it is a water-soluble chemotherapeutic agent that produces anti-cancer effects through the generation of DSBs [16]. In this regard, DOX and IR function similarly. While IR is used to treat adult GBM patients [17,18]. DOX is not due to its poor penetration through the blood-brain-barrier (BBB) [19]. However, significant progress has been made in developing prodrug analogs of DOX that can overcome this complication. For example, aldodoxorubicin is a promising prodrug of DOX that shows efficacy against GBM. At the chemical level, aldodoxorubicin contains a pH sensitive linker that allows for preferential release within GBM cells after efficient transport across the BBB [20]. Results from a recent phase II clinical trial with relapsed GBM patients showed that aldoxorubicin treatment correlated with positive tumor responses [21]. Since the mechanism of action for aldoxorubicin and DOX are identical, it is reasonable to predict that 5-NIdR will also potentiate the cell-killing effects of aldoxorubicin. The clinical value of this approach would be to eliminate drug-resistant cancer cells at low doses of the DNA-damaging agent, thereby reducing the risk of side effects [22].

Results from this study indeed show that combining 5-NIdR with DOX produces a synergistic increase in cell death in several GBM cell lines. This result was similarly reported when 5-NIdR is combined with TMZ [13]. However, quantitative analyses show that the cellular mechanisms accounting for cell death differ. In particular, combining 5-NIdR with DOX produces a classic apoptotic response involving caspase activation whereas combining 5-NIdR with TMZ induces cell death via mitotic catastrophe in a caspase-independent manner. This difference could reflect unique outcomes for inhibiting the activity of distinct DNA polymerases involved in replicating DNA lesions produced by DOX or TMZ.

## 2. Materials and Methods

### Cells and cell culture methods

U87, A172, MG118, and SW1088 cells were obtained from ATCC and cultured in a humidified atmosphere of 5% CO_2_ at 37 °C. Cells were grown in Dulbecco’s Modified Eagle Medium (DMEM) (Cellgro) supplemented with 10% fetal bovine serum (FBS) (Biowest), 1.0% Penicillin Streptomycin (Gibco) at 37 °C with 5.0% CO_2_. Since all cell lines were obtained from ATCC, informed consent was not required. Cell lines were routinely authenticated based on morphology and growth characteristics. All cells were expanded and then frozen at low passage (passages 2-5) within 2 weeks after the receipt of the original stocks. All cells used for experiments were between passages 6 and 12. Cell lines were tested for mycoplasma after each thaw or every 4 weeks when grown in culture. Mycoplasma infection was detected using the MycoAlert Mycoplasma Detection kit from Lonza (Walkersville, MD).

### Reagents

Phosphate-buffered saline (PBS), antibiotic and antifungal agents, amphotericin, propidium iodide, PrestoBlue, DAPI, Alexa Fluor 488, and apoptosis assay kit containing Alexa Fluor 488-labeled Annexin V were from Invitrogen. Doxorubicin was purchased from SigmaAldrich (>98% purity). Temozolomide was purchased from ChemPacific (>98% purity). 5-NIdR was synthesized and purified as previously described [22, 23].

### Cell viability assays

5-NIdR was added to wells in a dose-dependent manner (1−100 ΒιοAρχηιϖε g/mL). DOX was added to wells in a dose-dependent manner using concentrations ranging from 100 pM to 1 M. In all experiments, the final concentration of the co-solvent, DMSO, was maintained at 0.1%. In all cases, cells were treated for variable time periods for 72 hr. Cell viability was assessed using a Muse Cell Count (EMD Millipore) and via Cell-Titer blue assays. IC_50_ values for 5-NIdR and DOX were obtained using a non-linear regression curve fit of the data to Equation 1.

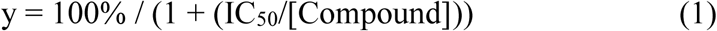

LD_50_ values for the artificial nucleoside and DNA damaging agent were calculated using identical approaches.

### Apoptosis Measurements

Cells were plated at an initial density of 200,000 cells/mL. Cells were treated with fixed concentrations of DMSO, 5-NIdR (100 μg /mL), DOX (100 nM), or a combination of 5-NIdR and DOX. At variable time intervals (24-72 hours), media containing DOX and/or 5-NIdR was removed and washed with 1X PBS. Cells were then treated with 0.25% trypsin, harvested by centrifugation, washed in PBS, and re-suspended in 100 μL of binding buffer containing 5 μM of Annexin V-Alexa Fluor 488 conjugate. Cells were treated with 1 μg/ mL PI and incubated at room temperature for 15 min followed by flow cytometry analysis. Cells were analyzed using a Muse Cell analyzer. 15,000-gated events were observed for each sample.

### Reactive Oxygen Species Measurements

U87 cells were plated at an initial density of 200,000 cells/mL and then treated with fixed concentrations of DMSO, 5-NIdR (100 μg/mL), DOX (100 nM), or a combination of 5-NIdR and DOX. At variable time intervals (24-72 hours), media containing DOX and/or 5-NIdR was removed and washed with 1X PBS. Cells were then treated with 0.25% trypsin, harvested by centrifugation, washed in PBS, and re-suspended in1X Assay Buffer supplied by the manufacturer (Luminex) at cell density of 10^6^ cells/mL 10 µL of suspended cells were added to a microfuge tube containing 190 µL of Muse Oxidative Stress Reagent working solution. The samples were mixed thoroughly by pipetting up and down for 5 seconds and then incubated at 37°C for 30 minutes. Cells were analyzed using a Muse Cell analyzer. 15,000-gated events were observed for each sample.

### Cell Cycle Analyses

Cells were plated at an initial density of 200,000/ml and treated with a fixed concentrations of DMSO, 5-NIdR (100 μg/mL), DOX (100 nM), or a combination of 100 μg/mL 5-NIdR and 100 nM DOX. Cell cycle analysis on fixed, permeabilized cells was performed using propidium iodide (PI) (Invitrogen). Approximately 10^6^ cells were fixed in 500 μL cold methanol and stored at 4 °C overnight. Cells were re-suspended in 500 μl propidium iodide solution (1 μg/ml propidium iodide, 0.1% (v/v) Triton, 0.1% (w/v) NaN_3_, 1.2% (v/v) 100 μg/ml RNAse A) and incubated for 1 hour at 37 °C. Cells were analyzed using either Muse Cell analyzer or Beckman Coulter EPICS-XL with EXPO 32 Data Acquisition software. 15,000-gated events were observed for each sample.

### Measurement of Caspase Activation

U87 cells were plated at an initial density of 200,000/ml overnights and then treated with a fixed concentrations of DMSO, 5-NIdR (100 μg/mL), DOX (100 nM), or a combination of 100 μg/mL 5-NIdR and 100 nM DOX. After 72 hours, media containing DOX and/or 5-NIdR was removed and replaced with fresh media. Cells were then treated with 0.25% trypsin, harvested by centrifugation, washed in PBS, and re-suspended in 1X Assay Buffer (50 μL per 100,000 cells). 5 µL of Muse® MultiCaspase Reagent working solution was added to a microfuge tube containing cells obtained from various treatments with DMSO, doxorubicin, 5-NIdR, the combination of doxorubicin and 5-NIdR, temozolomide, and temozolomide and 5-NIdR. The solution was mixed thoroughly by vortexing at medium speed for 5 seconds. Samples were incubated for 30 minutes in the 37°C incubator with 5% CO_2_. After incubation, 150 µL of Muse Caspase 7-AAD working solution was added to each tube and mixed thoroughly by vortexing at medium speed for 5 seconds. Samples were then Incubated at room temperature for 5 minutes in the dark. Cells were analyzed using either Muse Cell analyzer. 15,000-gated events were observed for each sample.

### Measurement of H2AX Activation

U87 cells were plated at an initial density of 200,000/ml overnights and then treated with fixed concentrations of DMSO, 5-NIdR (100 μg/mL), DOX (100 nM), or a combination of 100 μg/mL 5-NIdR and 100 nM DOX. After 72 hours, media containing DOX and/or 5-NIdR was removed and replaced with fresh media. Cells were then treated with 0.25% trypsin, harvested by centrifugation, washed in PBS, and re-suspended in 1X Assay Buffer (50 μL per 100,000 cells). Cells were permeabilized by adding ice-cold 1X permeabilization buffer and incubated on ice for 10 minutes. Cells were again centrifuged at 300 x g for 5 minutes, re-suspended in 1X assay buffer Cells were resuspended in 500 μL of 1X Assay Buffer per one million cells. Equal parts of Fixation Buffer were added to cell suspension (1:1) and mixed by gente pipetting up and down. Cells were incubated on ice for 5 minutes and then centrifuged at 300 x g for 5 minutes in a tabletop centrifuge. After discarding the supernatant, cells were permeabilized by adding 1 mL ice-cold 1X Permeabilization Buffer per one million cells. Cells were incubated on ice for 5 minutes and then centrifuged at 300 x g for 5 minutes in a tabletop centrifuge. After discarding the supernatant, cells were resuspended in 450 μL 1X Assay Buffer per one million cells. 90 μL of the supernatant was transferred to a fresh microfuge tube. 10 μL of the antibody working cocktail solution was added to each microcentrifuge tube containing the cell suspension. Cells were incubated for 30 minutes in the dark at room temperature. Following this incubation step, 100 μL of 1X Assay Buffer was to each microcentrifuge testing sample and centrifuged at 300 x g for 5 minutes on a tabletop centrifuge. The supernatant was discarded and cells were resuspended with 200 μL of 1X Assay Buffer. Cells were analyzed using a Muse Cell analyzer. 15,000-gated events were observed for each sample.

### Statistical analyses

The significance of difference in the mean value was determined using a two-tailed Student’s t-test and normal distribution was assumed in all cases. A one-way ANOVA analysis was used to compare the effects of cells treated with the combination of 5-NIdR and DOX versus treatment with DMSO, 5-NIdR, and DOX alone to determine p-values. P-values <0.05 were considered significant. All calculations were performed using KaleidaGraph software. All cell culture experiments were reproduced at least three times and performed on separate occasions independently. The number of samples and replicates for each experiment are provided in the text and figure legend.

## Results

### Potentiating the Cell Killing Effects of Doxorubicin

Initial studies examined the potency of DOX against U87 cells by measuring cell viability as a function of increasing concentrations of the DNA damaging agent over a 3-day period. Figure 2A provides a dose-response curve quantifying cell viability as a function of increasing concentrations of DOX (100 pM to 1 μM). A fit of the data to equation 1 yields an LD_50_ value of 250 +/- 30 nM. Based on this analysis, a sub-lethal concentration (<LD_50_) of 100 nM DOX was employed in all subsequent experiments. It was previously demonstrated that 5-NIdR produces minimal cytostatic and cytotoxic effects at concentrations up to 100 μg/mL [13]. These results were re-verified using flow cytometry to quantify cell viability and the onset of apoptosis using Annevin V staining coupled with PI uptake. Figure 2B provides dose-response curves using both assays on U87 cells treated with concentrations of 5-NIdR ranging from 10 to 100 μg/mL over a 3-day period. As indicated, treatment with the highest concentration of 5-NIdR (100 μg/mL) caused less than 10% cell death. Based on these results, a sub-lethal concentration of 100 μg/mL 5-NIdR was used in all subsequent experiments.

**Figure 2.**
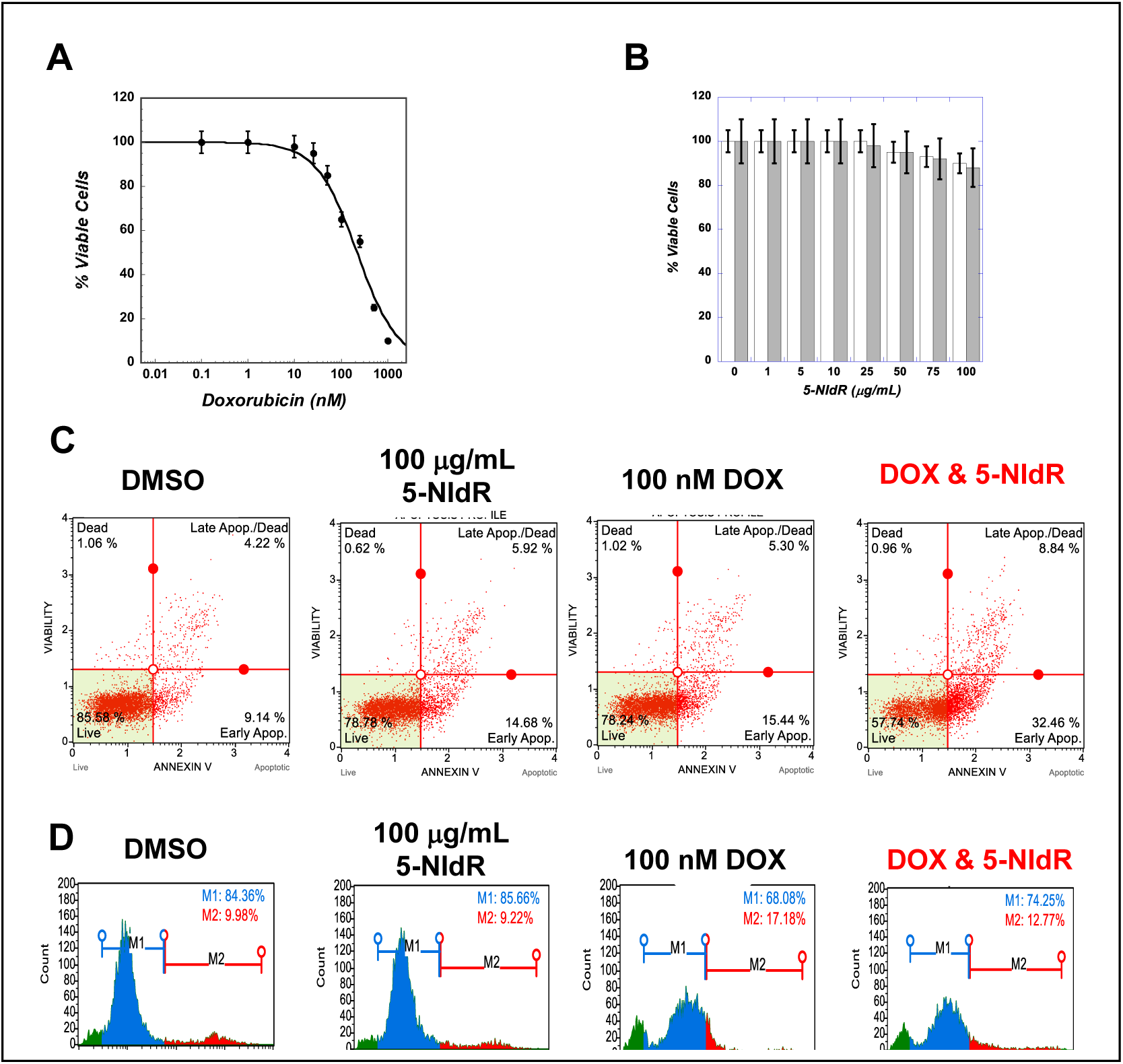
(A) Dose-response curve measuring the anti-cancer effects of doxorubicin (DOX) yields an LD_50_ value of 250 +/- 30 nM. (B) Dose-response curve measuring the anti-cancer effects of 5-NIdR. (C) Representative flow cytometry plots using PI uptake and annexin V staining to define live, early and late stage apoptotic, and necrotic cells as a function of treatment with DMSO, 5-NIdR, DOX, and a combination of DOX and 5-NIdR. (D) Representative histograms showing the fraction of reactive oxygen species (ROS) positive and negative U87 cells treated with DOX alone and combined with 5-NIdR. ROS negative cells are represented in blue while ROS positive cells are represented in red.

The next goal was to evaluate if treating U87 cells with a sub-lethal dose of 5-NIdR (100 μg/mL) could increase the efficacy of a sub-lethal dose of DOX (100 nM). Representative histograms using PI uptake and annexin V staining provided in Figure 2C show that U87 cells co-treated with 5-NIdR and DOX have higher levels of early and late stage apoptosis compared to cells treated with DOX or 5-NIdR alone. Table 1 summarizes results from of an average of four (4) independent experiments. Close inspection shows that the net apoptotic effect (Δ = 27.9%) for combining 5-NIdR with DOX is ∼2-fold greater than the additive effects for DOX and 5-NIdR treatment (Δ = 14.6%).

**Table 1.** Summary of dual parameter flow cytometry measuring apoptosis in U87 cells. Values represent an average of four (4) independent determinations performed on different days. Values in parenthesis represent the difference in percent early and late stage apoptosis in cells treated with each agant alone or combined as compared to treatment with DMSO (vehicle control).

| Condition | Viable | Early Apoptotic | Late Apoptotic | Necrotic | Total Apoptotic |
| --- | --- | --- | --- | --- | --- |
| DMSO | $86.8 \pm 2.5\%$ | $8.0 \pm 0.5\%$ | $4.1 \pm 0.7\%$ | $1.1 \pm 0.2\%$ | $12.1 \pm 0.8\%$<br>(0%) |
| 100 nM DOX | $81.2 \pm 3.5\%$ | $11.1 \pm 0.9\%$ | $5.5 \pm 0.7\%$ | $2.2 \pm 0.3\%$ | $16.6 \pm 0.9\%$<br>(4.5%) |
| 100 $\mu\text{g/mL}$<br>5-NIdR | $78.4 \pm 2.2\%$ | $12.9 \pm 1.1\%$ | $6.9 \pm 0.8\%$ | $1.8 \pm 0.2\%$ | $19.8 \pm 0.8\%$<br>(7.7%) |
| Combination | $59.6 \pm 1.8\%$ | $28.9 \pm 1.9\%$ | $9.7 \pm 1.0\%$ | $1.8 \pm 0.3\%$ | $38.6 \pm 1.3\%$<br>(26.5%) |

### Reactive Oxygen Species Generated by Doxorubicin

In addition to producing DSBs, DOX also generates reactive oxygen species (ROS) which can promote cell death by modifying cellular proteins and lipids [24,25]. Thus, the ability of 5-NIdR to enhance the cytotoxic effects of DOX could result from increasing ROS production rather than by inhibiting TLS activity. To assess this possibility, cellular levels of ROS production caused by DOX treatment were quantified in the absence and presence of 5-NIdR. Figure 2D shows that treatment with 100 μg/mL 5-NIdR has no effect on the population of ROS-positive cells compared to treatment with DMSO. In contrast, treatment with 100 nM DOX increases the percentage of ROS positive cells compared to cells treated with DMSO. However, U87 cells co-treated with 5-NIdR and DOX display nearly the same percentage of ROS-positive cells as cells treated with DOX alone. These data suggest that the increase in cell death produced by combining 5-NIdR with DOX is not caused by higher levels of ROS production.

### Induction of Apoptosis by Caspase Activation

Caspases are a family of proteases that play key roles in various forms of programmed cell death including apoptosis, pyroptosis, and necroptosis [26]. The Muse® MultiCaspase Assay was used to quantify the activity of several initiator and executioner caspases (caspase-1, 3, 4, 5, 6, 7, 8, and 9, respectively) in U87 cells treated with DOX in the absence and presence of 5-NIdR. Representative histograms provided in Figure 3A show that cells co-treated with 5-NIdR and DOX display higher levels of caspase activation compared to cells treated with DOX or 5-NIdR independently. Results summarized in Table 2 show that DMSO treatment produces low levels of both viable/caspase positive (2.9%) and non-viable/caspase positive (6.8%) cells. Likewise, U87 cells treated with 100 μg/mL 5-NIdR show low levels of viable/caspase positive (5.5%) and non-viable/caspase positive (8.1%) cells. This negative result is important as the lack of caspase activation suggests that the artificial nucleoside does not act as an ATP surrogate to activate the apoptosome [27]. Treatment with 100 nM DOX produces more pronounced effects as the percentage of viable/caspase positive and non-viable/caspase positive cells increase to 8.7% and 11.7%, respectively. Thus, the additive effect of treatment with 5-NIdR and DOX is a 14.8% net increase in total caspase activation (Table 2). However, combining 5-NIdR with DOX leads to significantly higher amounts of caspase activation, particularly with respect to the viable/caspase positive cell population (24.9%). In this case, combining 5-NIdR with DOX produces a ∼2-fold net increase in caspase activation compared to DOX and 5-NIdR used independently (compare 26.3% versus 14.8%, respectively). These results demonstrate that 5-NIdR increases the cytotoxicity of DOX by activating the intrinsic apoptotic pathway.

**Figure 3.**
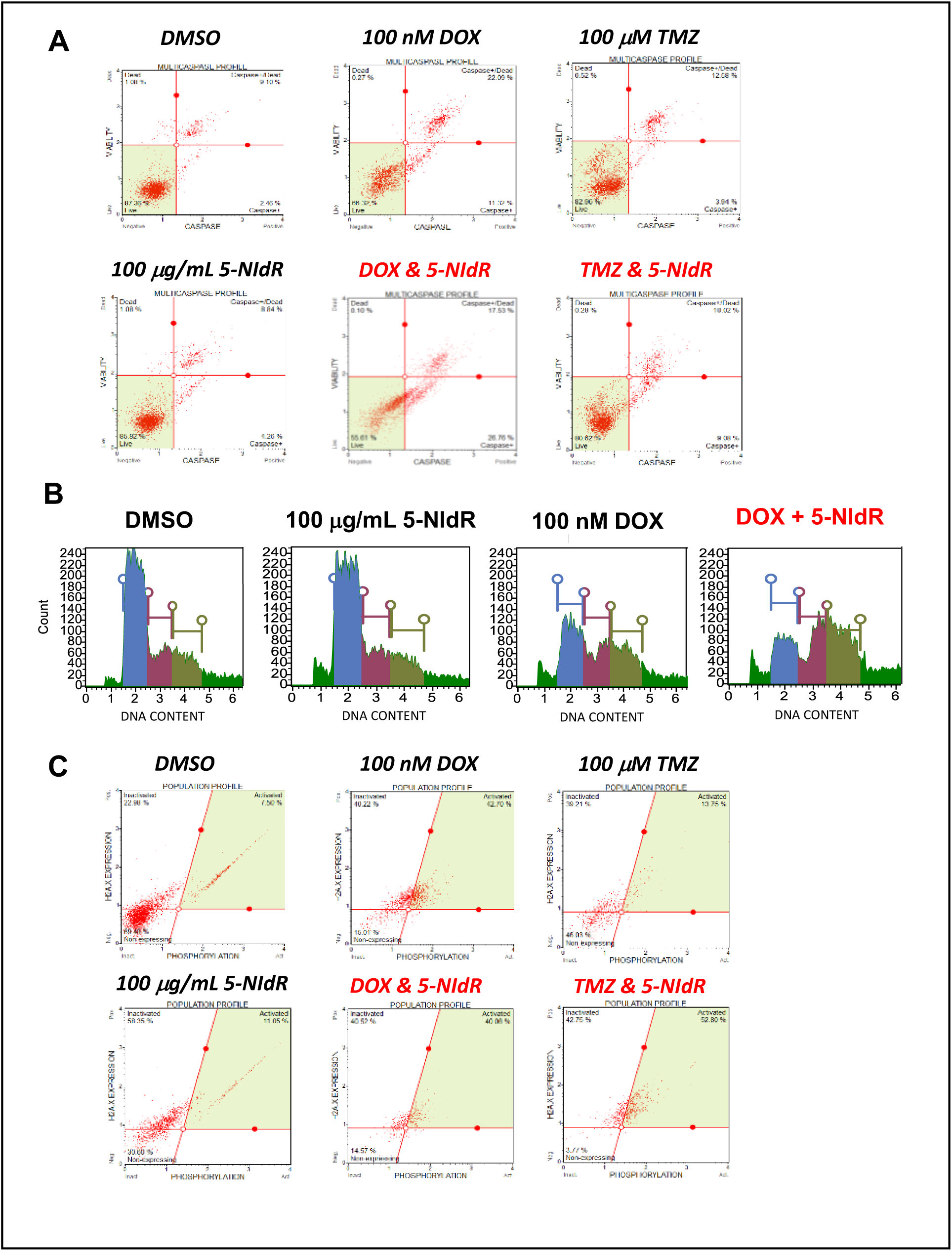
(A) Representative flow cytometry plots identifying caspase activation in cell treated with DMSO, 5-NIdR, DOX, a combination of DOX and 5-NIdR, TMZ, and a combination of TMZ and 5-NIdR. Co-treatment of U87 cells with 5-NIdR and DOX causes a synergistic activation of caspase activation while the combination of 5-NIdR and TMZ produces an additive effect in caspase activation. (B) Cell-cycle analyses for U87 cells treated with doxorubicin in the absence and presence of 5-NIdR. Combining 5-NIdR with DOX blocks cell-cycle progression at G_2_/M and likely reflects the ability of the analog to inhibit DNA polymerases involved in processing DSBs generated by DOX. (C) Flow cytometry histograms quantifying H2AX activation in U87 cells treated with DOX in the absence and presence of 5-NIdR. While DOX treatment increases the percentage of cells containing active γH2AX, the combination of DOX and 5-NIdR DOX does not significantly alter the distribution of cells expressing H2AX or the distribution of cells with active γH2AX. In contrast, the combination of 5-NIdR with TMZ significantly increases the percentage of cells expressing H2AX and the percentage of active γH2AX.

**Table 2.** Summary of caspase activation caused by treating U87 cells with DOX and TMZ alone and in the presence of 5-NIdR. Values represent an average of five (5) independent determinations performed on different days. Values in parenthesis represent the difference in percent caspase activation in cells treated with 5-NIdR, DOX, the combination of 5-NIdR and DOX, TMZ, and the combination of 5-NIdR and TMZ as compared to treatment with DMSO (vehicle control).

| Condition | Viable | Caspase /<br>Viable | Caspase<br>Non-Viable | Non-Viable | Total<br>Caspase |
| --- | --- | --- | --- | --- | --- |
| DMSO | $89.6 \pm 2.0\%$ | $2.9 \pm 0.3\%$ | $6.8 \pm 1.2\%$ | $0.7 \pm 0.1\%$ | $9.8 \pm 0.8\%$<br>(0%) |
| 100 $\mu\text{g/mL}$<br>5-NIdR | $85.6 \pm 2.6\%$ | $5.5 \pm 0.6\%$ | $8.1 \pm 1.5\%$ | $0.8 \pm 0.3\%$ | $13.6 \pm 0.9\%$<br>(3.8%) |
| 100 nM DOX | $78.7 \pm 1.5\%$ | $8.7 \pm 0.8\%$ | $11.7 \pm 1.1\%$ | $0.5 \pm 0.1\%$ | $20.8 \pm 1.2\%$<br>(11.0%) |
| DOX &<br>5-NIdR | $63.0 \pm 1.4\%$ | $24.9 \pm 1.7\%$ | $11.2 \pm 1.0\%$ | $0.3 \pm 0.1\%$ | $36.1 \pm 0.9\%$<br>(26.3%) |
| 100 $\mu\text{M}$ TMZ | $82.8 \pm 2.9\%$ | $5.7 \pm 0.9\%$ | $9.8 \pm 0.9\%$ | $0.4 \pm 0.1\%$ | $15.5 \pm 0.9\%$<br>(5.7%) |
| TMZ &<br>5-NIdR | $82.7 \pm 3.1\%$ | $8.6 \pm 1.2\%$ | $9.4 \pm 1.1\%$ | $0.3 \pm 0.1\%$ | $19.4 \pm 1.6\%$<br>(9.6%) |

The effect of combining 5-NIdR with DOX on caspase activation was compared to treatment with TMZ in the absence and presence of 5-NIdR. As summarized in Table 2, treatment with 100 μM TMZ alone produces small effects on both viable/caspase positive (5.7%) and non-viable/caspase positive cells (9.8%). When compared to vehicle-treated cells, the net effect of TMZ treatment on caspase activation is 5.7%. Thus, the net additive effect of TMZ and 5-NIdR is 9.5% (3.8% for 5-NIdR + 5.7% for TMZ). Surprisingly, the combination of 5-NIdR and TMZ produces only a 9.6% net increase in total caspase activation, an amount equal to the additive effects of TMZ and 5-NIdR independently (compare 9.6% versus 9.5%, respectively). These results directly contrast those observed for combining 5-NIdR with DOX which activates the intrinsic apoptotic pathway. Collectively, these results suggest that 5-NIdR reinforces a classic apoptotic response when combined with DOX but induces mitotic catastrophe when combined with TMZ.

### Cell-Cycle Analyses

The effect of combining 5-NIdR with DOX on cell cycle progression quantitatively assessed if the increase in caspase activation is caused by perturbations in cell-cycle progression. Figure 3B provides representative histograms for various treatments while Table 3 summarizes the results of ten independent determinations. In all experiments, a baseline for cell-cycle progression was first quantified by treating cells with DMSO for three days. Cells treated with DMSO display a pattern consistent with an asynchronous cell population in which the majority of cells exist at G_0_/G_1_ (52%) while smaller populations exist at S-phase (18%), G_2_/M (16%), and sub-G_1_ (13%). As previously reported, treatment with 5-NIdR has little effect on cell cycle progression as populations at G_0_/G_1_ (50%), S-phase (10%), G_2_/M (17%), and sub-G1 DNA (14%) are nearly identical to those compared DMSO treated cells. This result again indicates that the artificial nucleoside is relatively inert when used in the absence of an exogenous DNA damaging agent [12, 13]. In contrast, treatment with 100 nM DOX significantly affects cell cycle progression as G_0_/G_1_ levels are reduced (20%) which coincide with an increase in the population of cells at S-phase (25%). More pronounced effects are observed with increases in G_2_/M (31%) and sub-G1 DNA (24%). In general, these effects are consistent with the ability of DOX to produce both cytostatic and cytotoxic effects by creating DSBs. However, combining 5-NIdR with DOX produces some unexpected effects on cell-cycle progression. In particular, this combination causes a slight increase in G_0_/G_1_ (33%) and S-phase (28%) populations while decreasing the population of cells at G_2_/M (24%) compared to treatment with DOX alone. Perhaps the most surprising result is the observation of low levels of sub-G1 DNA (14%) that are essentially identical to those measured with DMSO treatment. The effect of increasing G_0_/G_1_ could result from the ability the artificial nucleotide to interfere with DSB repair which blocks cell cycle progression.

**Table 3.**
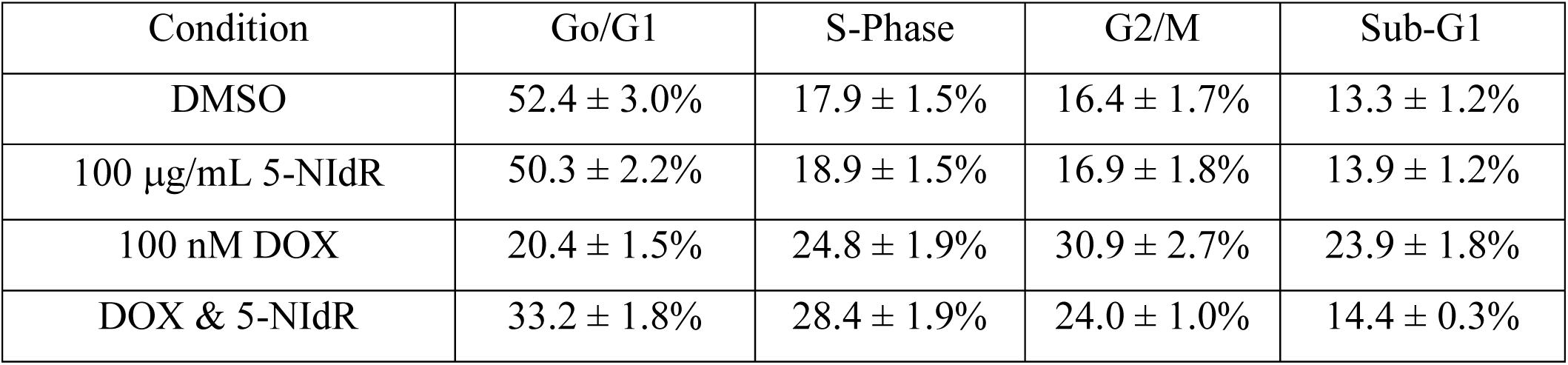
Summary of cell cycle analysis of U87 cells treated with DOX in the absence and presence of 5-NIdR. Values represent an average of ten (10) independent determinations performed on different days.

| Condition | Go/G1 | S-Phase | G2/M | Sub-G1 |
| --- | --- | --- | --- | --- |
| DMSO | 52.4 ± 3.0% | 17.9 ± 1.5% | 16.4 ± 1.7% | 13.3 ± 1.2% |
| 100 µg/mL 5-NIdR | 50.3 ± 2.2% | 18.9 ± 1.5% | 16.9 ± 1.8% | 13.9 ± 1.2% |
| 100 nM DOX | 20.4 ± 1.5% | 24.8 ± 1.9% | 30.9 ± 2.7% | 23.9 ± 1.8% |
| DOX & 5-NIdR | 33.2 ± 1.8% | 28.4 ± 1.9% | 24.0 ± 1.0% | 14.4 ± 0.3% |

### Examining the effects of DOX and 5-NIdR on H2AX Activation

DSB formation activates H2AX through phosphorylation of Ser139 to form γH2AX [28, 29]. Thus, quantifying γH2AX formation provides a specific and sensitive molecular marker to quantify if 5-NIdR affects the formation and subsequent repair of DSBs generated by DOX. Figure 3C provides representative results examining γH2AX formation in response to DOX treatment in the absence and presence of 5-NIdR. Table 4 summarizes the results from three (3) independent determinations. U87 cells treated with DMSO display a bimodal distribution of H2AX expression in which 28% of cells do not express H2AX while 72% express the protein. In the sub-population of cells expressing H2AX, only 10% is activated (γH2AX) while the remaining 62% remains inactive (H2AX). Treatment with 100 μg/mL 5-NIdR slightly shifts the distribution of H2AX expressing cells (65%) versus non-expressing (35%) cells. However, there is no change in the percentage of cells possessing γH2AX (10%), indicating that 5-NIdR treatment alone does not influence DSB formation. In contrast, treatment with 100 nM DOX strongly shifts H2AX expressing (93%) versus non-expressing (7%) cells. In addition, DOX treatment produces a large increase in the percentage of cells containing active γH2AX (∼50%). Both effects are consistent with the ability of DOX to generate DSBs. Surprisingly, combining 5-NIdR with DOX does not significantly affect the distribution of cells expressing H2AX (9%) or those with active γH2AX (47%). This result suggests that 5-NIdR does not increase the number of DSBs produced by DOX. However, it cannot eliminate the possibility that 5-NIdR actually inhibits the repair of DSBs produced by DOX.

**Table 4.** Summary of H2AX activation in U87 cells. Values represent an average of three (3) independent determinations performed on different days. Values in parenthesis represent the difference in percent caspase activation in cells treated with 5-NIdR, DOX, the combination of 5-NIdR and DOX, TMZ, and the combination of 5-NIdR and TMZ as compared to treatment with DMSO (vehicle control).

| Condition | Non-expressing cells | Cells expressing H2AX |  |
| --- | --- | --- | --- |
|  |  | Inactive | Active |
| DMSO | 28.2 ± 3.3% | 62.0 ± 5.2% | 9.8 ± 2.5% |
| 100 µg/mL 5-NIdR | 34.6 ± 3.5% | 55.1 ± 4.2% | 10.3 ± 3.0% |
| 100 nM DOX | 6.9 ± 2.1% | 43.4 ± 3.5% | 49.7 ± 3.9% |
| DOX & 5-NIdR | 8.8 ± 1.3% | 44.0 ± 3.8% | 47.2 ± 4.2% |
| 100 µM TMZ | 44.4 ± 2.5% | 23.6 ± 2.2% | 32.0 ± 2.9% |
| TMZ & 5-NIdR | 17.5 ± 1.9% | 33.1 ± 2.8% | 49.5 ± 3.5% |

H2AX activation in response to TMZ treatment in the absence and presence of 5-NIdR was also examined (Table 4). Treatment with 100 μM TMZ produces a bimodal distribution in H2AX expression with 44% of cells classified as non-expressing and 56% expressing H2AX. In the population expressing H2AX, 32% is activated (γH2AX) while the remaining 24% is inactive (H2AX). The higher levels of γH2AX likely reflect the formation of DSBs caused by the ability of DNA adducts to transiently disrupt the continuity of DNA replication. Consistent with this mechanism, combining 5-NIdR with TMZ produces a significant shift in the distribution of U87 cells expressing H2AX (82%) versus those that do not express H2AX (18%). More importantly, combining the two agents significantly increases the percentage of active γH2AX (∼50%). This result strongly suggests that 5-NIdR inhibits the activity of DNA polymerases that replicate DNA lesions formed by TMZ treatment. This inhibition disrupts the continuity of leading and lagging strand DNA synthesis to ultimately increase DSB formation.

### Efficacy in Other Astrocytoma Cell Lines

The next goal was to test if combining 5-NIdR with DOX produces similar cell killing effects in grade 3 (SW1088) and grade 4 (A172 and MG118) astrocytomas. A multiplexed approach described above was employed to quantify the effects of combining 5-NIdR with DOX on cell viability, apoptosis, and cell cycle progression across multiple cell lines. Results summarized in Table 5 show that the A172 cell line responds similarly to U87 cells when 5-NIdR is combined with sub-lethal concentrations of DOX or TMZ. While the magnitude of the increase in early and late-stage apoptosis is lower compared to U87 cells, it is clear that 5-NIdR potentiates the cell-killing effects of both drugs in A172 cells. Furthermore, similar effects are observed on cell-cycle progression and caspase activation in response to combining 5-NIdR with DOX.

**Table 5.** Summary of dual parameter flow cytometry measuring apoptosis, cell-cycle effects, and caspase activation in grade 3 and grade 4 astrocytoma cells after treatment with the combination of TMZ and 5-NIdR versus the combination of DOX and 5-NIdR.

| Cell Line<br>(Drug Combination) | Apoptosis | Cell Cycle Effects | Caspase<br>Activation | Mutations |
| --- | --- | --- | --- | --- |
| U87<br>(5-NIdR & TMZ) | 4.2-fold<br>increase | Block at G2/M | 1.01-fold<br>increase | CDKN2A,<br>PTEN,<br>CDK2C |
| U87<br>(5-NIdR & DOX) | 2.2-fold<br>increase | Block at G2/M | 1.80-fold<br>increase | CDKN2A,<br>PTEN,<br>CDK2C |
| A172<br>(5-NIdR & TMZ) | 1.7-fold<br>increase | Block at S-Phase<br>and G2/M | 1.61-fold<br>increase | CDKN2A,<br>PTEN |
| A172<br>(5-NIdR & DOX) | 1.7-fold<br>increase | Block at S-Phase<br>and G2/M | 3.3-fold<br>increase | CDKN2A,<br>PTEN |
| U118MG<br>(5-NIdR & TMZ) | 1.1-fold<br>increase | Minimal effect | 1.20-fold<br>increase | CDKN2A,<br>PTEN, TP53 |
| U118MG<br>(5-NIdR & DOX) | 1.2-fold<br>increase | Minimal effect | 1.05-fold<br>increase | CDKN2A,<br>PTEN, TP53 |
| SW1088<br>(5-NIdR & TMZ) | 1.1-fold<br>increase | Minimal effect | 1.08-fold<br>increase | CDKN2A,<br>PTEN, TP53 |
| SW1088<br>(5-NIdR & DOX) | 1.1-fold<br>increase | Minimal effect | 1.01-fold<br>increase | CDKN2A,<br>PTEN, TP53 |

Surprisingly, U118MG and SW1088 cell lines respond differently when 5-NIdR is combined with either DOX and TMZ. In these cases, the effect on early and late-stage apoptosis is significantly lower than that measured against U87 and A172 cells. Likewise, there is no increase in caspase activation in response to combining 5-NIdR with DOX or TMZ. Finally, combining 5-NIdR with either drug does not significantly block cell-cycle progression at S-phase as was observed with U87 cells. At face value, these differences clearly indicate that combining 5-NIdR with TMZ or DOX does not generate universal effects against astrocytomas. While the underlying mechanism for this difference is not currently understood, it is likely that genetic differences among these cell lines are responsible. For example, while all four cell lines possess mutations in CDKN2A and PTEN, two cell lines (U118MG and SW1088) possess mutations in p53 whereas U87 and A172 cells are both p53 proficient (30). It is interesting that the p53-proficient cell lines display similar synergistic cell-killing effects in response to combining 5-NIdR with DOX or TMZ whereas the p53-deficient cell line do not. A provocative hypothesis is that p53 activity is intimately involved in coordinating TLS activity. Further efforts are underway to examine this possibility.

## Discussion

This study quantitatively assessed how brain cancer cells respond to DNA-damaging agents. A key feature was assessing the ability of 5-NIdR to potentiate the cytotoxic effects of TMZ and DOX and interrogate the role of TLS as a cellular mechanism driving drug resistance. While TMZ and DOIX create non-instructional DNA lesions (abasic sites versus DSBs, respectively), each lesion is processed by different DNA polymerases the repair or tolerance of the generated DNA damage. Table 6 highlights key differences in how GBM cells respond toward the combination of either drug with 5-NIdR. Based on these differences, a working model for the cellular responses to each DNA-damaging agent is provided in Figure 4. Previous studies demonstrated that 5-NIdR potentiates the cell killing effects of TMZ by ∼4-fold [12, 13]. While 5-NIdR also potentiates the cell-killing effects of DOX, the magnitude of this effect is ∼2-fold lower. Another important difference is the underlying mechanism responsible for cell death. Co-treatment with DOX and 5-NIdR generates an ∼2-fold increase in caspase activation whereas combining 5-NIdR with TMZ has no observable effect on caspase activity. This difference suggests that 5-NIdR reinforces a classic apoptotic response caused by DOX while the combination of 5-NIdR and TMZ induces cell death by mitotic catastrophe. Consistent with this model, cell-cycle experiments show differences in the responses to either drug combination. For example, combining 5-NIdR with TMZ causes an increase in sub-G_1_ levels of DNA by blocking replication during S-phase. In this model, 5-NIdR inhibit TLS activity causing cells to undergo cell death via mitotic catastrophe. In contrast, combining 5-NIdR with DOX blocks cell-cycle progression primarily at G_o_/G_1_, and this likely reflects the ability of the analog to inhibit one or more DNA polymerases that process DSBs generated by DOX.

**Figure 4.**
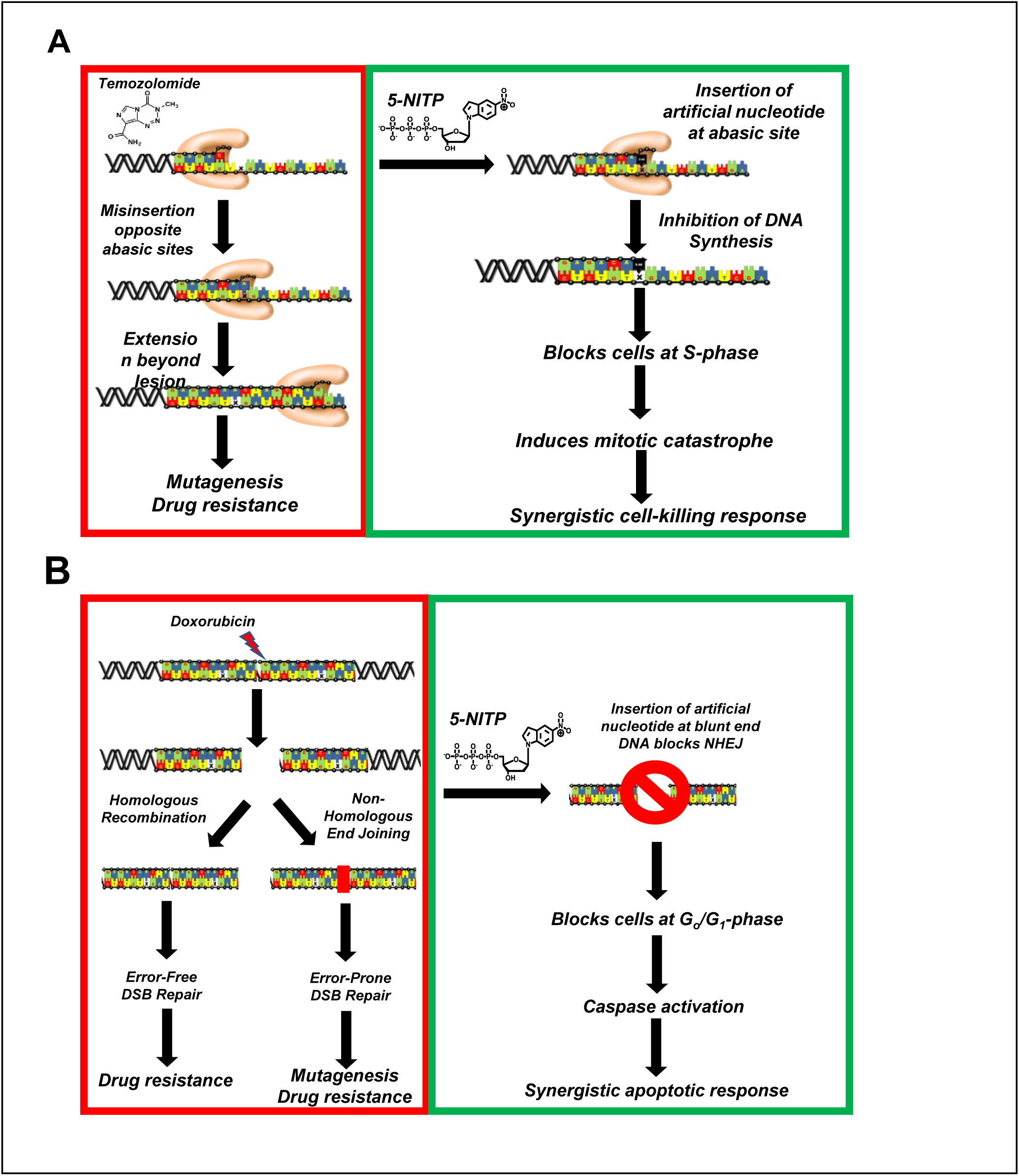
Hypothetical models accounting for the cellular differences in the ability of 5-NIdR to potentiate the cell killing effects of temozolomide and doxorubicin. (A) Temozolomide produces abasic sites that are misreplicated by several DNA polymerases. This activity generates resistance to temozolomide and increases the risk of mutagenesis. Since 5-NITP is preferentially incorporated opposite abasic sites, it terminates translesion DNA synthesis to induces cell death by mitotic catastrophe. (B) Doxorubicin produces double strand DNA breaks that are also non-instructional DNA lesions. 5-NITP may be used by one or more DNA polymerases involved in either homolgous recombination or non-homologous end joining. By inhibiting these polymerases, the artificial nucleotide induces apoptosis by inhibiting the timely repair of double strand DNA breaks.

**Table 6.**
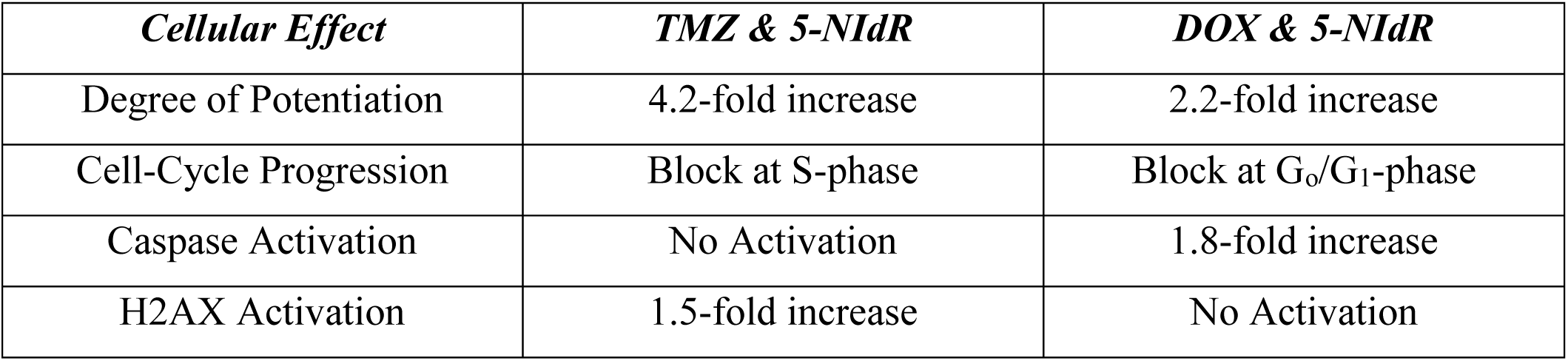
Comparison of the cellular phenotypes observed after treating U87 cells with the combination of TMZ and 5-NIdR versus the combination of DOX and 5-NIdR.

| *Cellular Effect* | *TMZ & 5-NIdR* | *DOX & 5-NIdR* |
| --- | --- | --- |
| Degree of Potentiation | 4.2-fold increase | 2.2-fold increase |
| Cell-Cycle Progression | Block at S-phase | Block at G<sub>0</sub>/G<sub>1</sub>-phase |
| Caspase Activation | No Activation | 1.8-fold increase |
| H2AX Activation | 1.5-fold increase | No Activation |

The next question is which DNA polymerase(s) is (are) inhibited by the artificial nucleotide to generate these differential responses. Previous *in vitro* studies demonstrated that high-fidelity DNA polymerases (pol ε and pol δ) and specialized DNA polymerases (pol η and pol ι) insert 5-NITP opposite an abasic site 100-fold more efficiently compared to dATP [13]. However, the specialized DNA polymerase, pol κ, and the repair DNA polymerases, pol λ and pol μ do not incorporate 5-NITP or dATP to an appreciable extent opposite an abasic site. Thus, the ability of 5-NIdR to potentiate TMZ likely reflects utilization of the corresponding nucleoside triphosphate, 5-NITP, by multiple DNA polymerases (pol ε, pol δ, pol η, and pol ι) which then blocks TLS. In the case of DOX, however, the underlying mechanisms for this potentiation are likely more complex due to the multiple pathways that can process DSBs. These include homologous recombination (HR), non-homologous end joining (NHEJ), and micro-homology mediated end joining (MMEJ) [31–35]. HR is typically an error-free repair pathway that utilizes high-fidelity DNA polymerases such as pol ε and pol δ [36] to replicate undamaged templates. The template-dependent nature of HR makes it unlikely that 5-NIdR directly affects this pathway since 5-NITP is not efficiently utilized by high-fidelity polymerases when replicating undamaged DNA [11]. In contrast, NHEJ is an error-prone pathway since the ends of DSBs are ligated without extensive homology between the two sequences [37, 38]. Two specialized DNA polymerase participate in this classic form of NHEJ. The first, pol λ, resynthesizes missing nucleotides at the break by binding both ends of a DNA break through extensive contacts with the 5’ phosphate of the downstream DNA strand [39, 40]. The second polymerase, pol μ, also resynthesizes damaged or missing nucleotides at the blunt end of DNA during NHEJ [41]. While it was previously demonstrated that pol λ and pol μ did not appreciably incorporate 5-NITP opposite an abasic site [13], it has not been determined if either DNA polymerase can incorporate the artificial nucleotide opposite blunt end DNA to influence DSB repair. This may be possible since the enhanced base-stacking properties of 5-NITP allows the artificial nucleotide to be efficiently incorporated by various DNA polymerases at blunt end DNA [42]. Finally, another specialized DNA polymerase, pol q, participates in MMEJ [43]. During this process, pol q uses homologous nucleotides from both ends of the DSB to initiate repair. Since this results in the loss of nucleotides in the DNA sequence, MMEJ is also highly mutagenic. Current efforts are underway to determine if pol q utilizes 5-NITP during blunt-end DNA synthesis.

## Conclusion

DNA-damaging agents such as doxorubicin play integral roles in treatment strategies for patients suffering from hematological and solid cancers. Unfortunately, resistance to these agents can develop, thereby reducing their overall efficacy. This report provides additional evidence for the use of artificial deoxynucleoside, 5-NIdR, that inhibits the inappropriate misreplication of DNA lesions produced by these agents. In particular, co-treatment of glioblastoma cells with doxorubicin and 5-NIdR leads to an increase in cancer cell death compared to treatment with either compound individually. This reflects an increase in apoptosis that appears independent of increasing reactive oxygen species that can produce adverse side effect. The co-treatment of cancer cells with 5-NIdR and DNA damaging agent may represent a new strategy to counteract drug resistance caused by inadequate DNA repair coupled with translesion DNA synthesis.

## Author Contributions

Conceptualization, Anthony Berdis; methodology, Anthony Berdis.; validation, Anthony Berdis.; formal analysis, Anthony Berdis.; investigation, Anthony Berdis.; data curation, Anthony Berdis.; writing—original draft preparation, Anthony Berdis.; writing—review and editing, Anthony Berdis.; visualization, Anthony Berdis.; supervision, project administration, Anthony Berdis.; funding acquisition, Anthony Berdis. All authors have read and agreed to the published version of the manuscript.”

